# Increasing the efficiency of long-read sequencing for hybrid assembly with k-mer-based multiplexing

**DOI:** 10.1101/680827

**Authors:** Alexander Dilthey, Sebastian A. Meyer, Achim J. Kaasch

## Abstract

Hybrid genome assembly has emerged as an important technique in bacterial genomics, but cost and labor requirements limit large-scale application. We present Ultraplexing, a method to improve per-sample sequencing cost and hands-on-time of Nanopore sequencing for hybrid assembly by at least 50%, compared to molecular barcoding while maintaining high assembly quality (Quality Value; QV ≥ 42). Ultraplexing requires the availability of Illumina data and uses inter-sample genetic variability to assign reads to isolates, which obviates the need for molecular barcoding. Thus, Ultraplexing can enable significant sequencing and labor cost reductions in large-scale bacterial genome projects.

## Background

Accurate characterization of large numbers of microbial genomes is becoming increasingly important in microbiology. For example, bacterial genome-wide association studies (bGWAS) rely on the sequencing of large numbers of samples to correlate genetic variants to phenotypes such as antibiotic resistance or virulence (1–3). Further examples are phylogenetic analyses and quality assurance in technical microbiology (4–7).

A variety of sequencing technologies with different technological trade-offs have emerged for the sequencing of microbial genomes. Short-read sequencing technologies (such as Illumina (8)) have low error rates (<0.1%) but provide only limited resolution of complex and repetitive genomic regions. An example are the genes encoding *S. aureus* protein A (*spa*) and fibronectin binding-protein (*fnbpA*), which play key roles in the pathogenesis of *S. aureus (9)* and which cannot be reliably assembled from short-read data (10). Long-read sequencing technologies (Pacific Biosciences (11), Oxford Nanopore (12, 13)) generate sequencing reads of tens or even hundreds of kilobases in length, enabling the correct structural resolution of complex regions; their higher error rates (5 – 15%), however, can negatively impact consensus and small-variant genotyping accuracy (14–16).

Combining short- and long-read data has therefore emerged as a standard approach for the resolution of bacterial genomes (17). Long-read sequence information can be used to deconvolute short-read-based assembly graphs (hybrid *de novo* assembly; (18–21)). Alternatively, *de novo* assemblies from long reads (22) can be polished with short-read data to improve consensus accuracy (23). By either approach, the coverage requirements to arrive at a high-quality assembly of a microbial genome are typically modest (50-100X for each data type; (24, 25)).

Molecular barcoding approaches enable the cost-effective sequencing of multiple samples in one run (“multiplexing”). Molecular barcoding involves the labeling of each DNA sample with a unique barcode sequence; pooling and joint sequencing of the samples; and determining the source sample for each sequencing read, based on its barcode sequences. Highly efficient, automated implementations of molecular barcoding exist for the Illumina platform, enabling the sequencing of hundreds of microbial isolates to sufficient coverage with a single flow cell. Molecular barcoding approaches for long-read platforms, however, are less effective. A maximum of 24 samples can currently be multiplexed on an Oxford Nanopore MinIon flow cell. In addition, the preparation of multiplex libraries requires significant hands-on time (>12h compared to 3h for a non-multiplexed library), comes with significant losses of input material, and, presumably, the pipetting steps reduce attainable read lengths by shearing. These factors make barcoded long-read sequencing costly and labor-intensive, and the availability of a more scalable approach to multiplexed long-read sequencing would be highly desirable.

Here we present Ultraplexing, a new method that allows the pooling of multiple samples in long-read sequencing without relying on molecular barcodes. Ultraplexing uses inter-sample genetic variability, as measured by Illumina sequencing, to assign long reads to individual isolates (Figure 1). Specifically, each isolate genome is represented by its de Bruijn graph, constructed from sample-specific short-read data; and each long read is assigned to the sample de Bruijn graph it is most compatible with (or randomly in cases of a draw). A similar approach enables haplotype-aware assembly in eukaryotic genomes (26).

**Figure 1:**
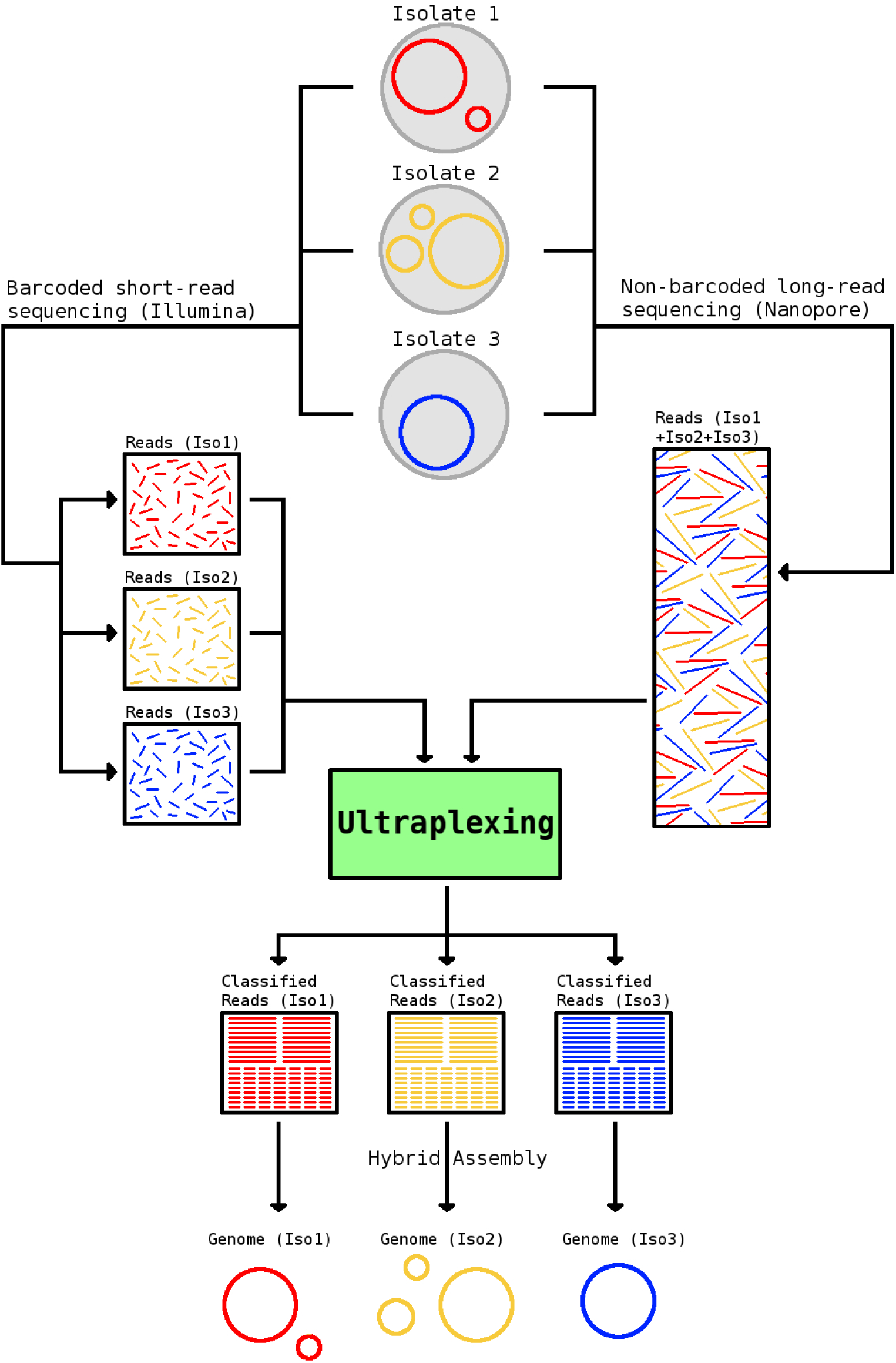
Overview of the Ultraplexing approach. Long reads are generated in simple pooled sequencing runs. The Ultraplexing algorithm determines the most likely source genome for each long read by carrying out a comparison between the read and the de Bruijn graphs of the sequenced sample genomes, inferred from short-read data. Hybrid assembly of sample-specific long and short reads enables the recovery of complete bacterial genomes.

The intuition behind Ultraplexing is that there will typically be a high-quality alignment between a read and the assembly graph of the source genome it emanates from. Importantly, the assignment of reads completely contained in genomic regions shared among multiple samples (e.g. due to mobile genetic elements or inter-sample genetic homology) may remain ambiguous. This, however, will typically have no or only a small effect on the accuracy of the hybrid assembly process, for the affected reads will spell equally valid assembly graph traversals in all compatible samples.

Ultraplexing requires the availability of Illumina data. It is applicable to studies that either incorporate the generation of these from the beginning or it can serve as a cost-effective method to generate additional long-read data for samples that have already been short-read sequenced. In the following, we demonstrate that Ultraplexing can match or even outperform classical molecular long-read barcoding approaches in terms of assembly quality while enabling significant reductions in cost and hands-on time.

## Results

We used simulated and real Nanopore and Illumina sequencing data (Supplementary Table 9) to evaluate the performance of Ultraplexing in the context of bacterial hybrid *de novo* assembly. In all experiments, we relied on Unicycler as an established method for hybrid assembly (18). We primarily focused on the quality of the generated assemblies, i.e. structural accuracy (number of contigs, reference recall, assembly precision) and consensus accuracy (Single Nucleotide Polymorphisms; SNPs), measured against the utilized reference genomes (in simulations) or barcoding-based assemblies (for real data). To distinguish between Ultraplexing-mediated effects and intrinsic assembly complexity for the selected isolates, we reported assembly accuracy for random (in all experiments) and perfect (in simulations) assignment of long reads. Additionally, we assessed the proportion of correctly assigned reads. Of note, all simulation experiments were based on conservative assumptions (e.g. 5 Gb throughput per long-read flow cell; see Methods for further details), and no mis-assemblies were identified through visual inspection in any of the Ultraplexing-based sets.

### Simulation experiment I: Multi-species Ultraplexing

In a first step, we evaluated Ultraplexing on a sample of 10 different clinically important bacterial species (Supplementary Table 1, Supplementary Figure 1). The Ultraplexing algorithm assigned all but 2 of 477,890 simulated long reads to the correct bacterial isolate (close to 100% classification accuracy, s. Supplementary Figure 1). Ultraplexing-based assemblies were highly concordant with the underlying reference genomes, achieving near-perfect structural agreement (average reference recall and assembly precision >99.999%) and low divergence (average number of SNPs against the reference genome: 57). Furthermore, assembly accuracy metrics for Ultraplexing and perfect read assignment were virtually identical (for example, an average of 57 SNPs for Ultraplexing compared to 56 SNPs for perfect assignment).

### Simulation experiment II: Single-species Ultraplexing with 10 – 50 isolates

To assess Ultraplexing performance on closely related isolates and with increasing sample numbers, we randomly selected sets of 10, 20, 30, 40, and 50 genomes from 181 publicly available complete assemblies of the human pathogen *Staphylococcus aureus* (Supplementary Table 5). Of note, as simulated long-read flow cell capacity was held constant, sets with more genomes contained less long-read data per isolate. Across experiments, the proportion of correctly assigned reads decreased as sample numbers increased and varied between 35% and 95% (Figure 2A). To test whether reduced read assignment accuracies were due to inter-sample sequence homologies, we computed the metric *Δedit distance* for random samples of mis-assigned reads and found an average *Δedit distance* of 0.3%, with more than 50% of mis-assigned reads exhibiting a *Δedit distance* of 0 (Figure 2B). At the read alignment level, the genomes that the mis-assigned reads were assigned to are thus indistinguishable or very similar to the true source genomes. Consistent with this, the generated Ultraplexing-based assemblies were highly concordant with the utilized reference genomes (average reference recall ≥ 99.96% and assembly precision ≥ 99.99% across sets; average number of SNPs 46; Figure 2C-F). Furthermore, assembly accuracy metrics for Ultraplexing and perfect read assignment were comparable even with increasing number of bacterial isolates; for example, the average number of SNPs per genome in the run with 50 bacterial isolates was 59 for Ultraplexing (QV 47) and 32 for perfect read assignment (QV 49). Complete results for this experiment are presented in Supplementary Table 2 and visualized in Figure 2.

**Figure 2:**
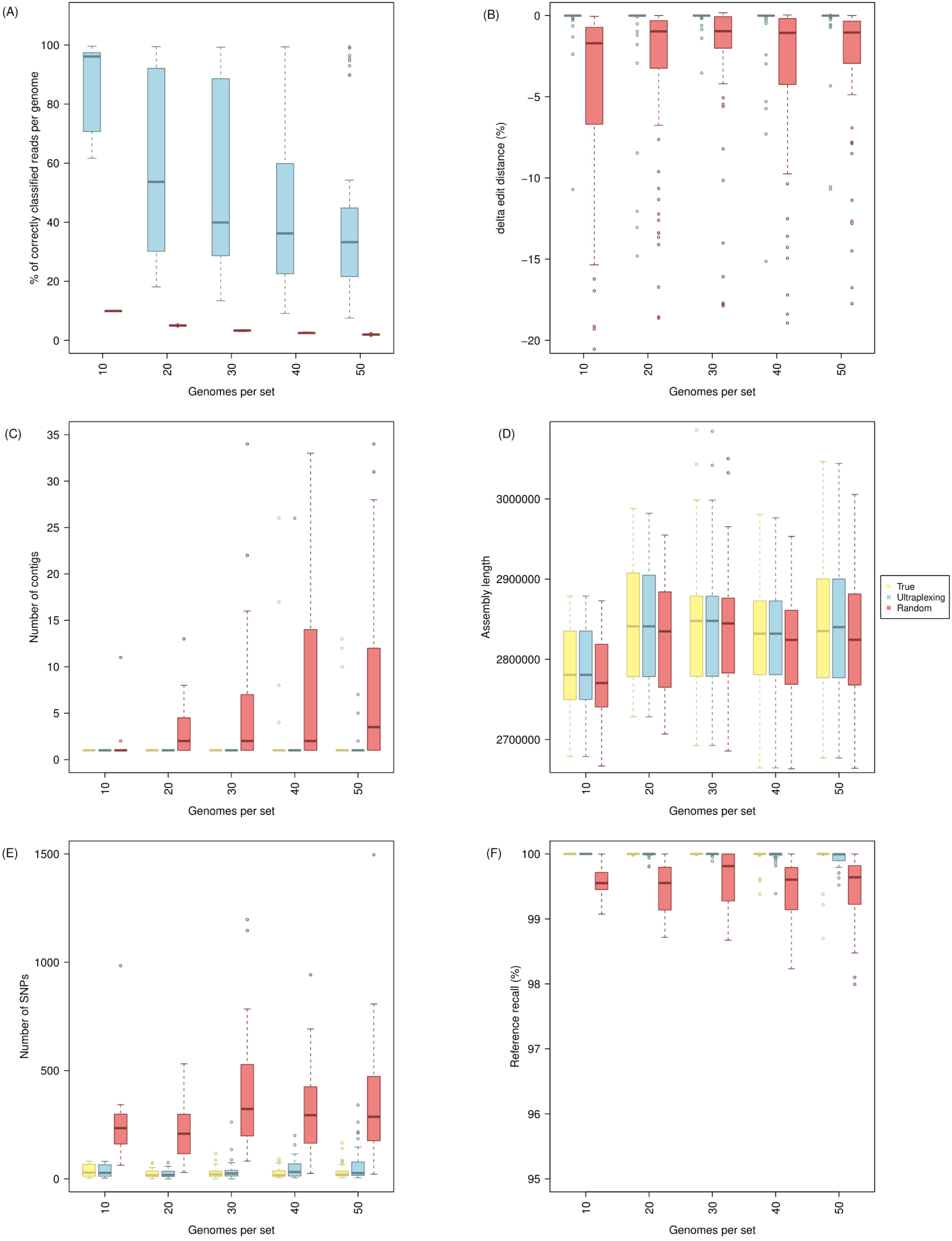
Simulated Ultraplexing runs with 10 – 50 *S. aureus* genomes, in comparison to perfect (True) and random (Random) assignment of long reads. The figure shows the proportion of correctly assigned long reads (A); Δedit distance for random samples of falsely classified long reads (B); the distribution of contigs per assembly (C); the distribution of assembly lengths (D); the distribution of SNPs per assembly (E); and the distribution of reference recall (F). SNPs and reference recall were calculated relative to the utilized reference genomes, and all metrics within the same set of genomes are based on the same simulated short-read data.

### Simulation experiment III: Impact of plasmids

In addition to the chromosomal genome, many bacterial cells harbor plasmids. Plasmids are extrachromosomal circular strings of DNA that are generally much smaller than the chromosomal DNA. Plasmids can vary in copy number within each cell and they often exhibit complex and repetitive sequence structures. Since plasmid sequences could reduce the performance of the Ultraplexing algorithm, we repeated the previous simulation experiments with sets of 10 - 50 *S. aureus* genomes that all harbored plasmids. We found that the accuracy of chromosomal genome assemblies was not affected by the presence of plasmids. Additionally, the plasmid recovery rate was comparable to assemblies based on reads assigned to their true source; complete recovery was achieved in 135 of 150 total isolate genomes with Ultraplexing, and in 137 with perfect read assignments. Identified reasons for incompletely recovered plasmids included high sequence homology to other plasmids or the genomic DNA (Supplementary Table 6). Complete results for this experiment are presented in Supplementary Table 3 and visualized in Supplementary Figure 4 (chromosomal genome) and Supplementary Figure 5 (plasmids). Possible reasons for incompletely recovered plasmids are high sequence homology to other plasmids or the genomic DNA and the consequential merging with these (Supplementary Table 6).).

### Real-data experiment I: Nanopore-based Ultraplexing of 10 *S. aureus* clinical samples

To assess the performance of Ultraplexing on real data, we randomly selected ten bacterial isolates of the species *Staphylococcus aureus* from our collection of clinical isolates. To generate a reference genome for each isolate, we sequenced each sample on an Illumina system, performed barcoded Oxford Nanopore sequencing with the 12-sample barcoding kit (∼214X coverage per isolate; mean read length 8.3 kb), and carried out hybrid *de novo* assembly. The generated reference genomes consist of 1 - 3 circular contig per isolate, representing the chromosomal genome (∼2.8 Mb in length) and plasmids (2.3 – 34.9 kb in length, all circular; BLAST (27) classification results are shown in Supplementary Table 7).

To test Ultraplexing on these isolates, we demultiplexed the barcoded Nanopore sequencing data with the Ultraplexing algorithm and carried out hybrid *de novo* assembly. The Ultraplexing-based assemblies showed a high degree of concordance (Figure 3) with the generated reference genomes in terms of contig number, assembly length, genome structure (average reference recall and assembly precision > 99.9%), and consensus accuracy (4 SNPs per isolate on average and 6 of 10 isolates with no detected SNPs). In contrast, assemblies based on random read assignment yielded lower-quality assemblies across all considered metrics (for example, 136 SNPs per genome; Figure 3D). Complete results for all genomes are presented in Supplementary Table 4 and visualized in Figure 3. Summary statistics of the Illumina and Nanopore sequencing runs can be found in Supplementary Table 9.

**Figure 3:**
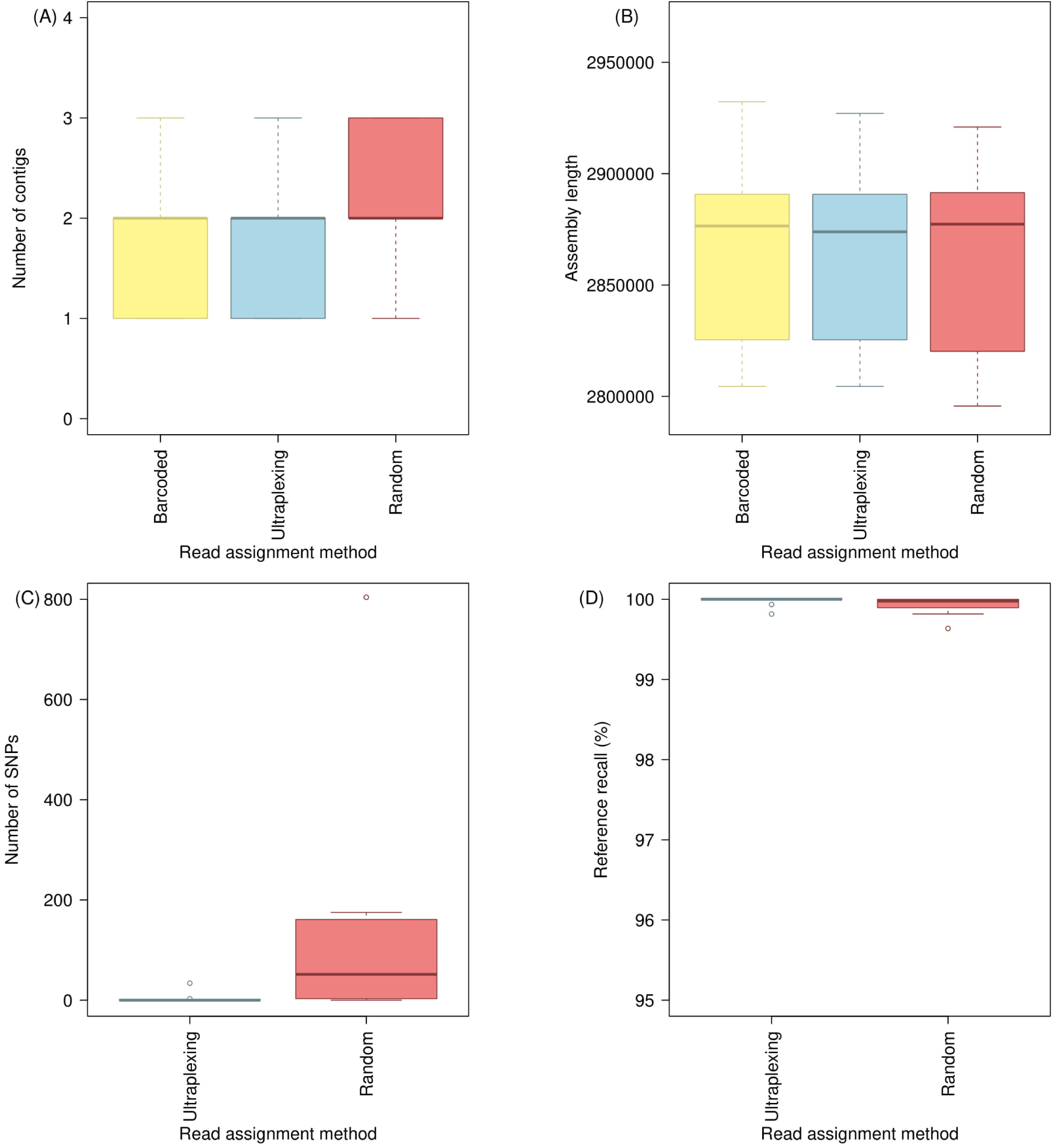
Ultraplexing and classical molecular barcoding on a set of ten *S. aureus* isolates. For different read assignment methods applied to the same set of Nanopore reads, the figure shows the distribution of contigs per assembly (A); the distribution of assembly lengths (B); the distribution of SNPs per assembly (C); and the distribution of reference recall (D). SNPs and reference recall were calculated relative to assemblies based on molecular barcoding, and the same Illumina sequencing data were used throughout. Barcoded: reads assigned according to molecular barcodes; Ultraplexing: reads assigned by the Ultraplexing algorithm; Random: reads assigned randomly.

### Read-data experiment II: Nanopore-based Ultraplexing of 48 clinical isolates

To assess the feasibility of applying Ultraplexing to a larger number of samples, we repeated the previous experiment with 48 samples. As in the previous experiment, barcoded Nanopore (∼235X coverage per isolate; average read length 5.6 kb) and Illumina (∼50X coverage per isolate; 2 × 250bp reads with MiSeq v2 chemistry) sequencing was carried out to generate reference genomes for the 48 samples. In contrast to the previous experiment, however, we employed a recently released 24-sample native barcoding Nanopore kit. We observed that the reference genomes so-generated exhibited significant variation in contig numbers (Figure 4A) and one outlier in terms of assembly size (Figure 4B).

**Figure 4:**
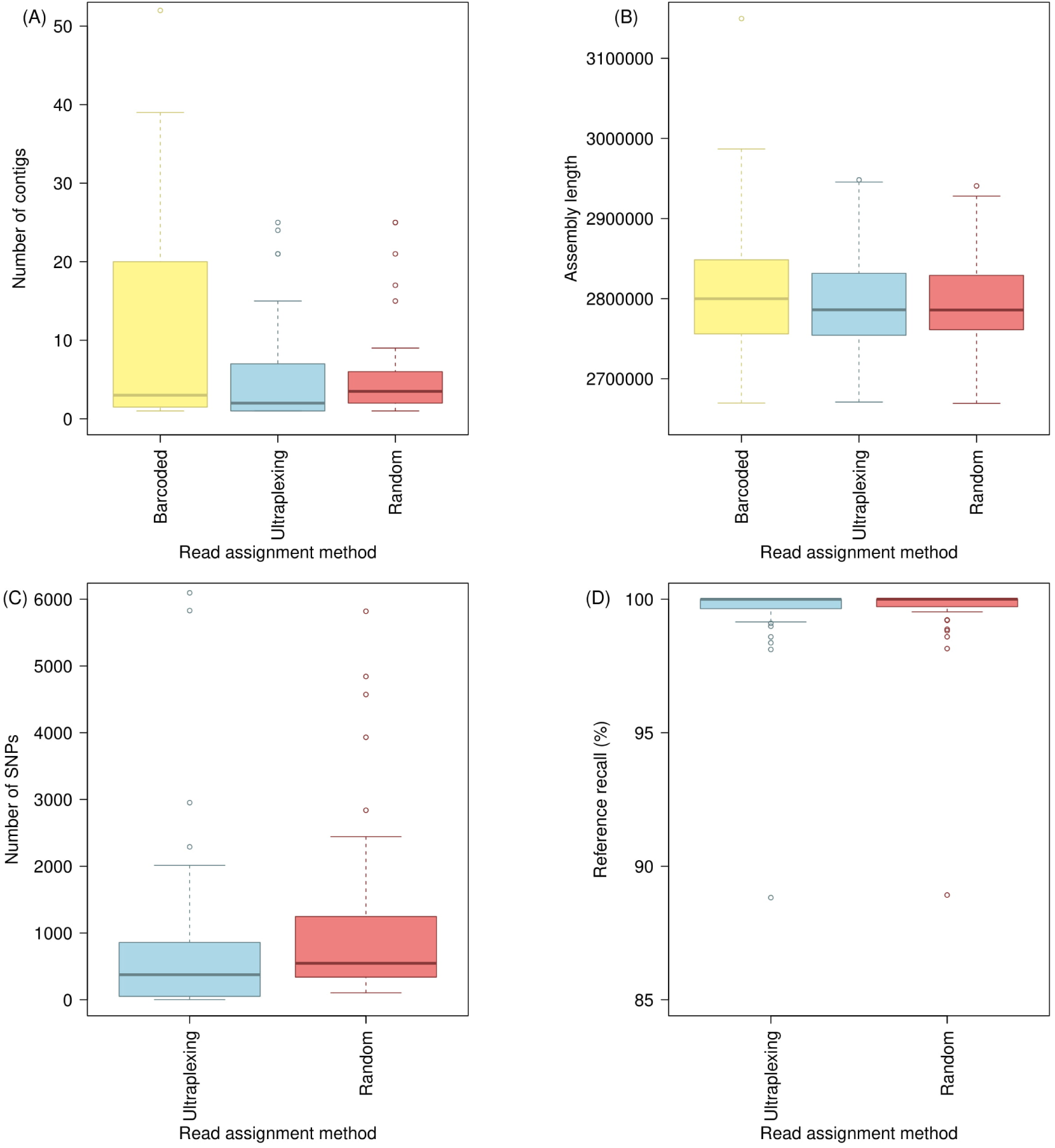
Ultraplexing and classical molecular barcoding on a set of 48 *S. aureus* isolates. The figure shows the distribution of contigs per assembly (A); the distribution of assembly lengths (B); the distribution of SNPs per assembly (C); and the distribution of reference recall (D). SNPs and reference recall are calculated relative to assemblies based on molecular barcoding, and the same Illumina sequencing data were used throughout. Barcoded: molecularly barcoded Nanopore data, 2 flow cells with 24 samples each; Ultraplexing: reads assigned by the Ultraplexing algorithm, 1 flow cell with 48 samples; Random: reads from the Ultraplexing run, assigned randomly.

For Ultraplexing, long-read sequencing data (∼95X coverage per isolate; average read length 11.7 kb) were generated in a single MinIon run by pooling DNA from the 48 isolates. Reads were demultiplexed with the Ultraplexing algorithm and hybrid *de novo* assembly was carried out. The generated assemblies exhibited a plausible profile in terms of contig numbers and assembly length (Figure 4). However, a comparison with the generated reference genomes showed a considerable degree of divergence. For example, we found an average of 784 SNPs between the Ultraplexing-based and barcoding-based assemblies (Figure 4C).

These findings prompted us to investigate whether the generated reference genomes were of sufficient quality. We therefore repeated the reference genome generation process and generated new short- and long-read sequencing data (166X and 384X coverage, respectively; average Nanopore read length 9.7 kb) for a set of 10 isolates with unusually large or fragmented reference assemblies, utilizing the same 12-sample barcoding kit as in the first real-data experiment. We repeated the hybrid *de novo* assembly process with the new data and obtained a set of 10 completely circularized genomes clustered around an average assembly length of 2.8 Mb (Figure 5). Using these as improved reference genomes for the selected 10 samples, we found a high degree of concordance between the Ultraplexing-based assemblies and the new reference genomes both in terms of genome structure (average reference recall and assembly precision >99.6%) and in the number of SNPs per genome (180 on average, equivalent to QV 42). Ultraplexing-based assemblies showed higher accuracy than the initial barcoding-based assemblies in terms of accuracy, which exhibited, for example, an average of 2,706 SNPs against the new reference genomes. Complete results for the comparison of the 48 Ultraplexing-based assemblies against the initial set of reference genomes are presented in Supplementary Table 4 and visualized in Figure 4; complete results for the selected 10 samples, comparing Ultraplexing-based assemblies and the initial reference genomes against the improved reference genomes, are shown in Supplementary Table 4 and visualized in Figure 5. Read length and coverage statistics for all sequencing runs can be found in Supplementary Table 9.

**Figure 5:**
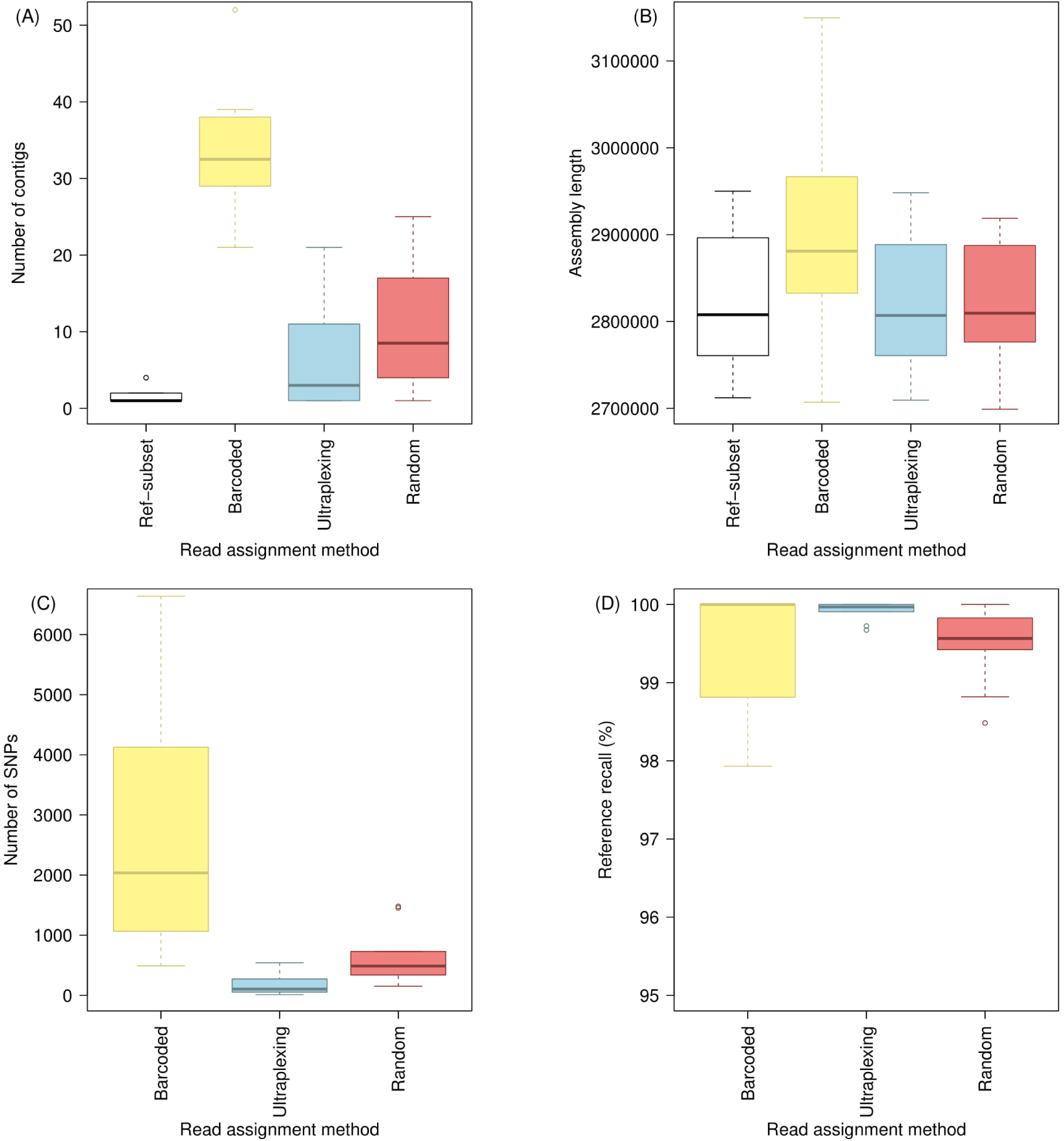
Ultraplexing, a classical molecular barcoding run, and a repeat molecular barcoding run on 10 of the 48 *S. aureus* isolates of the second real-data experiment. The figure shows the distribution of contigs per assembly (A); the distribution of assembly lengths (B); the distribution of SNPs per assembly (C); and the distribution of reference recall (D). The 10 isolates were selected because they exhibited implausibly large or fragmented assemblies in the first molecular barcoding run. The second molecular barcoding run (Ref-subset) was based on the 12-sample Nanopore barcoding kit, involved the generation of new Illumina and Nanopore data with more and (for Nanopore) longer reads, and yielded 10 high-quality assemblies; these were used as references against which SNPs and reference recall were calculated against (panels C and D). The data for Barcoded, Ultraplexing and Random are a subset of these visualized in Figure 4.

## Discussion

We have presented Ultraplexing, a method that resolves pooled long-read sequencing data in the context of hybrid *de novo* assembly without the use of barcoding. Ultraplexing leverages inter-sample genetic variation to assign pooled long reads to individual isolates and benefits from the fact that Illumina sequencing enables the reliable characterization of sample genome structure at the level of k-mers.

Using simulated sequencing data, we demonstrated that Ultraplexing enables the generation of highly accurate hybrid assemblies and reliably detects plasmids, even in datasets that contain multiple isolates of the same bacterial species. We have also validated the method on two real Nanopore sequencing datasets and shown that Ultraplexing-based assemblies are virtually identical to barcoding-based assemblies when comparing multiplexed runs with the same number of isolates. When using Ultraplexing to increase the number of samples over the current maximum of molecular barcoding approaches, Ultraplexing-based assemblies generally maintain high accuracy.

Of note, our results indicate that 48-sample Ultraplexing is closer in terms of assembly accuracy to molecular barcoding with 10 samples than to molecular barcoding with 24 samples. Furthermore, average consensus quality as reported in the 48-sample experiment (QV 42) likely represents an underestimate. This is because the selected set of repeat isolates was non-random with high assembly complexity, and because all differences between the Ultraplexing-based assemblies and the improved reference genomes were counted as errors in the former. Based on the first simulation experiment, however, in which an average of 50 SNPs per genome remained even with perfect read assignment, it seems likely that some of the detected differences in the 48-sample experiment represent errors in the utilized reference genomes. Thus, Ultraplexing is an accurate and cost-effective method for determining the genomes of large numbers of bacterial samples.

A key advantage of Ultraplexing in comparison to molecular barcoding is decreased cost and hands-on time. The number of samples sequenced per flow cell can at least be doubled and barcoding reagents are not necessary. Hands-on time was reduced eightfold in our 48-sample experiment (12 hours per flow cell with 24 barcoded samples compared to 3 hours for one Ultraplexing run with 48 samples). Taking into account potential differences in sample handling operator performance, we conservatively estimate that the hands-on-time benefit conferred by Ultraplexing is at least 50%. On the other hand, Ultraplexing can consume significant computational resources (70 CPU hours and 175Gb of memory for the demultiplexing step in the experiment with 48 samples).

The finding that Ultraplexing produced more accurate assemblies than conventional barcoding in a subset of samples despite lower overall coverage can likely be explained by read length. Reads in the Ultraplexing dataset of 48 isolates were significantly longer than in the barcoded samples (average read length of 11.67kb for Ultraplexing compared to 5.55kb for molecular barcoding; Supplementary Table 9), presumably facilitating hybrid assembly. These read length differences might be driven by pipetting-induced shearing during the barcoding protocol. Our results also indicate that the 12-sample barcoding kit might be preferable over the 24-sample barcoding kit on the Nanopore platform if the generation of reference-quality genomes with molecular barcoding kit is desired.

Although our primary focus was on assembly accuracy, we also evaluated the accuracy of individual read assignments in the simulation experiments. One important factor driving read assignment accuracy was the extent of genetic variability between the pooled samples. Consistent with this, Ultraplexing achieved near-perfect read assignment in the multi-species experiment but reduced assignment accuracy in the single-species experiment. We hypothesized that mis-assignments driven by inter-sample sequence homology would have a negligible effect on assembly accuracy. Consistent with this, assembly accuracy was relatively insensitive to increasing numbers of mis-assigned reads in the single-species experiment, and we could confirm that inter-sample sequence homology accounts for the majority of mis-assigned reads using edit distance metrics. Furthermore, assembly accuracy was significantly reduced for random read assignment, reflecting higher proportions of falsely assigned reads in the absence of underlying sequence homologies. The applicability of Ultraplexing in the context of a “long-read-first” approach (for example, Canu followed by Pilon) remains to be determined. In addition, Ultraplexing may be less well-suited for applications that depend on accurate assignments of individual reads, such as read-based methylation calling.

Our study has a number of limitations. First, we have only validated Ultraplexing on a single long-read technology, Oxford Nanopore. However, we expect Ultraplexing to work as well with the PacBio technology, based on its more random error profile (28, 29) and prior work demonstrating successful k-mer-based classification of eukaryotic PacBio reads (26). Second, we have focused on a limited set of clinically important bacterial species and not explored in depth how genome structure affects Ultraplexing. Thus, we cannot exclude the possibility that the performance of Ultraplexing may degrade when applied to certain bacterial species, e.g. due to large repeat structures in their genomes. Third, we have not rigorously tested the technical limits of Ultraplexing, including the maximum number of isolates and the necessary properties of the short-read sequencing data. Given that flow cell output has been increasing steadily, extraction of high-molecular weight DNA for long-read sequencing may plausibly become the most significant limiting factor. Fourth, in terms of bioinformatics methods development, Ultraplexing relies on simple k-mer statistics instead of proper graph alignment (30–32), and we have not explored methods for the optimization of intra-batch genetic diversity in large sequencing projects. These points could be addressed in future work.

## Conclusion

Ultraplexing is a new method for multiplexed long-read sequencing in the context of hybrid *de novo* assembly. Ultraplexing-based assemblies are highly accurate in terms of genome structure and consensus accuracy and exhibit quality characteristics comparable to assemblies based on molecular barcoding. Through increasing the number of samples per flow cell and simplified library preparation, Ultraplexing enables significant reductions of long-read sequencing costs and hands-on time. Thus, Ultraplexing enables the cost-effective complete resolution of large numbers of bacterial genomes.

## Methods

### The Ultraplexing read assignment algorithm

Let *n* denote the number of sequenced bacterial samples. We assume the availability of high-coverage Illumina sequencing data for each of the *n* individual isolates and that a pool of high-molecular-weight DNA, representing a mixture of the genomes of the *n* isolates, has been sequenced with a long-read sequencing technology like Oxford Nanopore or Pacific Biosciences. For each sample, a de Bruijn graph (k = 31) is constructed from the sample-specific Illumina data and the graph is cleaned (removal of low-coverage supernodes) with Cortex (17). Each long read from the pooled run is assigned to the sample for which the number of read k-mers present in the sample de Bruijn graph is maximal (or randomly in cases of a draw). We note that our approach can be understood as a heuristic approach to read-to-graph alignment.

### Hybrid assembly and assembly evaluation criteria

Unicycler (14) was used for all hybrid assembly experiments in this publication. For Ultraplexing, sample-specific long-read sequencing data were obtained by applying the Ultraplexing algorithm.

The performance of Ultraplexing was assessed (I) by assessing the proportion of reads assigned to the correct sample (in simulations); (II) by comparing the generated Ultraplexing-based hybrid *de novo* assemblies to reference genomes (downloaded from RefSeq for simulations and based on barcoding-based hybrid assembly for real data, see below); (III) by comparing the accuracy of Ultraplexing-based assemblies to that of assemblies based on random (all experiments) or perfect (in simulations) assignment of long reads.

To assess the accuracy of an assembly, we compared the assembly to the corresponding reference genome. As baseline characteristics, we considered the total number of contigs and the combined assembly length. Furthermore, nucmer v3.1 (33) was used to generate an alignment between the assembly and the reference genome, globally filtering identified diagonals with “delta-filter −1”. We used the filtered diagonals to compute three quality metrics: “SNPs”, measuring consensus accuracy; “reference recall”, the fraction of the reference covered by the assembly; “assembly precision”, the fraction of the assembly covered by the reference. When reported, QV is calculated as 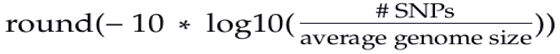 (Phred scale). Of note, assembly precision was close to 100% in all experiments, and we don’t separately report on this metric.

For the simulation experiment with plasmids, we separately evaluated the sets of chromosomal and plasmid contigs for each assembly. We relied on RefSeq annotations for determining the status (chromosomal or plasmid) of each contig in the reference and assigned the status of each assembly contig according to the status of the its highest-scoring nucmer hit in the reference.

### Read assignment accuracy and edit distance

In simulated datasets, we calculated the proportion of correctly assigned long reads. A read was counted as correctly assigned if, and only if, it was assigned to the genome it was simulated from. For mis-assigned reads, we additionally defined a metric referred to as “Δedit distance”, using edlib (34). Let d_1_ be the ends-free edit distance between a read and the genome it was simulated from and let d_2_ be the edit distance between a read and the genome it was assigned to. Δedit distance is defined as d_1 –_ d_2_, divided by the length of the read. A negative value indicates a better alignment to the source genome than to the predicted genome. To assess the distributional properties of Δedit distance, the metric was calculated for random samples of 100 mis-assigned reads per method.

### Simulation experiments

For the multi-species simulation experiments, chromosomal sequences of 10 clinically important species were downloaded from RefSeq (35). For the single-species experiments without plasmids, chromosomal sequences of 181 complete *S. aureus* genomes were downloaded from RefSeq (35). For the single-species simulation experiment with plasmids, 169 complete genomes were downloaded that contained between 2 and 11 annotated plasmids. The accessions of all downloaded genomes are listed in Supplementary Table 5, and the selected genome subsets are listed in Supplementary Tables 2 and 3.

For each genome, 300 Mb of short-read data were simulated with wgsim (version 0.3.1-r13)(36), using the parameters base error rate (-e 0.005), length of first read (−1 150), length of second read (−2 150), outer distance between the read ends (-d 278), standard deviation (-s 128), mutation rate (-r 0) and fraction of indels (-R 0). Long-read data were simulated with pbsim (version 1.0.3)(37), using the parameters prefix of the output (--prefix [prefix]), coverage (--depth 200), mean read-length (--length-mean 8370), standard deviation of the read-length (--length-sd 6389), maximum read-length (--length-max 61011), minimum read-length (--length-min 230), mean sequencing accuracy (--accuracy-mean 0.88) and model of quality code (--model_qc model_qc_clr). For all experiments, we assumed a constant long-read flow cell capacity of 5Gb, and per-isolate coverage was adjusted accordingly (i.e. 5Gb total output divided by the number of simulated isolates). Simulated long-read data were pooled and demultiplexed with the Ultraplexing algorithm. Hybrid *de novo* assembly was carried out and the generated assemblies were benchmarked against the utilized reference genomes.

### DNA extraction and sequencing

DNA was extracted from overnight bacterial cultures in 3 ml LB broth. For short read sequencing, the “DNeasy UltraClean Microbial” Kit was used according to the manufacturer’s instruction. 1 ng of DNA per isolate was used for the library preparation with the TruePrep DNA Library Prep Kit. Short-read sequencing was conducted on a MiSeq instrument (Illumina) using 250 bp paired end sequencing using v2 chemistry. DNA extraction for long-read sequencing was performed with the MagAttract HMW DNA Kit (QIAGEN). Wide bore pipette tips were used to avoid shearing. Long-read sequencing was carried out on a MinIon device with FLO-MIN106 flow cells and the SQK-LSK108 (first real-data experiment and 48-sample run in the second real-data experiment) and SQK-LSK109 (repeat run of 10 isolates in the second real-data experiment) ligation sequencing kits. Of note, SQK-LSK109 involves reduced pipetting, possibly decreasing shearing. For barcoded long-read sequencing, samples were labeled with barcodes using the Oxford Nanopore ligation sequencing kits (EXP-NBD103 kit or EXP-NBD114 for 12 and 24 samples per run, respectively), and reads were demultiplexed with Albacore (version 2.1.3). For Ultraplexing, DNA from individual samples was pooled based on equal weight to yield a total of 700ng of DNA, and demultiplexing was carried out with the Ultraplexing algorithm. Summary statistics of all sequencing runs are presented in Supplementary Table 9.

### Real-data validation experiments

For all experiments with real data, we used hybrid assembly with Unicycler (18) to generate high-quality reference genomes for all isolates, combining molecularly barcoded short- and long-read data.

Molecular long-read barcoding was carried out using the 12-sample barcoding kit (EXP-NBD103) for the first real-data experiment and for the 10-sample repeat sequencing run in the second real-data experiment; initial long-read generation for the complete set of 48 samples in the second experiment was based on the 24-sample barcoding kit (EXP-NBD104). Barcoded Illumina sequencing runs were carried out for the multi-species experiment; for the complete set of 48 samples in the single-species experiment; and for the repeat set of 10 genomes in the single-species experiment. All sequencing runs are summarized in Supplementary Table 9.

### Plasmid identification

To check if smaller contigs in real-data experiments (Figures 3 and 5) represented plasmids, we used the online version of BLAST (27). All non-chromosomal contigs (assumed to be the longest contig in each assembly) were blasted against the nucleotide (nt) database, restricted to sequences that correspond to bacteria (taxid:2), and if the best hit was characterized as plasmid, had a high identity (≥90%), and a low e-value (0 or close to 0), we assumed that the contig represented a correctly assembled plasmid (Supplementary Table 7).

## Supporting information

Supplementary Table 1 (multi-species)

Supplementary Table 2 (single-species)

Supplementary Table 3 (single-species plus plasmids)

Supplementary Table 4 (single-species, real data)

Supplementary Table 5 (used genomes)

Supplementary Table 6 (wrong assembled plasmids)

Supplementary Table 7 (found plasmids)

Supplementary Table 8 (read comparison)

Supplementary Table 9 (sequencing metrics)

## List of abbreviations

SNP: Single Nucleotide Polymorphism
QV: Quality Value

## Declarations

### Ethics approval and consent to participate

Not applicable

### Consent for publication

Not applicable

### Availability of data and material

The datasets generated and analysed during the current study are available under the BioProject accession number PRJNA528186).

The source code of the Ultraplexing algorithm is available from GitHub: https://github.com/SebastianMeyer1989/UltraPlexer.

The Ultraplexing algorithm is made available under the MIT license and implemented in C++, Perl and R. Sequence-to-graph alignment depends on the Cortex package version 1.0.5.21 (17).

### Authors’ contributions

AD and AJK: study concept and design, data management, data analysis, data interpretation, and manuscript writing. S.M: data management, data analysis and data interpretation, manuscript writing

All authors have read and approved the final draft submitted.

## Acknowledgements

We thank Lisanna Hülse (Heinrich-Heine University Düsseldorf) for technical assistance regarding DNA extraction, library preparation, and nanopore sequencing. We thank Harald Seifert (University of Cologne) for providing the bacterial isolates. Computational support and infrastructure were provided by the “Centre for Information and Media Technology” (ZIM) at the Heinrich-Heine University Düsseldorf (Germany).

## References

1. Falush D. Bacterial genomics: Microbial GWAS coming of age. Nat Microbiol. 2016 May;1:16059.

2. Chen PE, Shapiro BJ. The advent of genome-wide association studies for bacteria. Curr Opin Microbiol. 2015 Jun 1;25:17–24.

3. Young BC, Earle SG, Soeng S, Sar P, Kumar V, Hor S, et al. *Panton-Valentine leucocidin* is the key determinant of *Staphylococcus aureus pyomyositis* in a bacterial GWAS. eLife. 2019 Feb 22;8.

4. Jagadeesan B, Gerner-Smidt P, Allard MW, Leuillet S, Winkler A, Xiao Y, et al. The use of next generation sequencing for improving food safety: Translation into practice. Food Microbiol. 2019 Jun;79:96–115.

5. Cocolin L, Mataragas M, Bourdichon F, Doulgeraki A, Pilet M-F, Jagadeesan B, et al. Next generation microbiological risk assessment meta-omics: The next need for integration. Int J Food Microbiol. 2018 Dec 20;287:10–7.

6. Diaz-Sanchez S, Hanning I, Pendleton S, D’Souza D. Next-generation sequencing: The future of molecular genetics in poultry production and food safety. Poult Sci. 2013 Feb 1;92:562–72.

7. Taboada EN, Graham MR, Carriço JA, Van Domselaar G. Food Safety in the Age of Next Generation Sequencing, Bioinformatics, and Open Data Access. Front Microbiol [Internet]. 2017 [cited 2019 Jun 18];8. Available from: https://www.frontiersin.org/articles/10.3389/fmicb.2017.00909/full

8. Nakamura K, Oshima T, Morimoto T, Ikeda S, Yoshikawa H, Shiwa Y, et al. Sequence-specific error profile of Illumina sequencers. Nucleic Acids Res. 2011 Jul;39:e90.

9. Menzies BE. The role of *fibronectin* binding proteins in the pathogenesis of *Staphylococcus aureus* infections. Curr Opin Infect Dis. 2003 Jun;16:225–9.

10. Bartels MD, Petersen A, Worning P, Nielsen JB, Larner-Svensson H, Johansen HK, et al. Comparing Whole-Genome Sequencing with Sanger Sequencing for spa Typing of *Methicillin*-Resistant *Staphylococcus aureus*. J Clin Microbiol. 2014 Dec;52:4305–8.

11. Rhoads A, Au KF. PacBio Sequencing and Its Applications. Genomics Proteomics Bioinformatics. 2015 Oct;13:278–89.

12. Mikheyev AS, Tin MMY. A first look at the Oxford Nanopore MinION sequencer. Mol Ecol Resour. 2014;14:1097–102.

13. Laver T, Harrison J, O’Neill PA, Moore K, Farbos A, Paszkiewicz K, et al. Assessing the performance of the Oxford Nanopore Technologies MinION. Biomol Detect Quantif. 2015 Mar 24;3:1–8.

14. Krishnakumar R, Sinha A, Bird SW, Jayamohan H, Edwards HS, Schoeniger JS, et al. Systematic and stochastic influences on the performance of the MinION nanopore sequencer across a range of nucleotide bias. Sci Rep [Internet]. 2018 Feb 16 [cited 2019 Mar 14];8. Available from: https://www.ncbi.nlm.nih.gov/pmc/articles/PMC5816649/

15. Utturkar SM, Klingeman DM, Land ML, Schadt CW, Doktycz MJ, Pelletier DA, et al. Evaluation and validation of de novo and hybrid assembly techniques to derive high-quality genome sequences. Bioinformatics. 2014 Oct;30:2709–16.

16. Jain M, Koren S, Miga KH, Quick J, Rand AC, Sasani TA, et al. Nanopore sequencing and assembly of a human genome with ultra-long reads. Nat Biotechnol. 2018 Apr;36:338–45.

17. Iqbal Z, Caccamo M, Turner I, Flicek P, McVean G. De novo assembly and genotyping of variants using colored de Bruijn graphs. Nat Genet. 2012 Jan 8;44:226–32.

18. Wick RR, Judd LM, Gorrie CL, Holt KE. Unicycler: Resolving bacterial genome assemblies from short and long sequencing reads. PLoS Comput Biol [Internet]. 2017 Jun 8 [cited 2019 Mar 14;13(6). Available from: https://www.ncbi.nlm.nih.gov/pmc/articles/PMC5481147/

19. Bankevich A, Nurk S, Antipov D, Gurevich AA, Dvorkin M, Kulikov AS, et al. SPAdes: A New Genome Assembly Algorithm and Its Applications to Single-Cell Sequencing. J Comput Biol. 2012 May;19:455–77.

20. Antipov D, Korobeynikov A, McLean JS, Pevzner PA. hybridSPAdes: an algorithm for hybrid assembly of short and long reads. Bioinformatics. 2016 Apr 1;32:1009–15.

21. Zerbino DR, Birney E. Velvet: Algorithms for de novo short read assembly using de Bruijn graphs. Genome Res. 2008 May;18:821–9.

22. Koren S, Walenz BP, Berlin K, Miller JR, Bergman NH, Phillippy AM. Canu: scalable and accurate long-read assembly via adaptive k-mer weighting and repeat separation. Genome Res. 2017 May;27:722–36.

23. Walker BJ, Abeel T, Shea T, Priest M, Abouelliel A, Sakthikumar S, et al. Pilon: An Integrated Tool for Comprehensive Microbial Variant Detection and Genome Assembly Improvement. PLoS ONE [Internet]. 2014 Nov 19 [cited 2019 Mar 19];9(11). Available from: https://www.ncbi.nlm.nih.gov/pmc/articles/PMC4237348/

24. Koren S, Harhay GP, Smith TP, Bono JL, Harhay DM, Mcvey SD, et al. Reducing assembly complexity of microbial genomes with single-molecule sequencing. Genome Biol. 2013;14:R101.

25. Wick RR, Judd LM, Gorrie CL, Holt KE. Completing bacterial genome assemblies with multiplex MinION sequencing. Microb Genomics [Internet]. 2017 Sep 14 [cited 2019 Mar 19;3(10). Available from: https://www.ncbi.nlm.nih.gov/pmc/articles/PMC5695209/

26. Koren S, Rhie A, Walenz BP, Dilthey AT, Bickhart DM, Kingan SB, et al. De novo assembly of haplotype-resolved genomes with trio binning. Nat Biotechnol. 2018 Dec;36:1174–82.

27. Altschul SF, Gish W, Miller W, Myers EW, Lipman DJ. Basic local alignment search tool. J Mol Biol. 1990 Oct 5;215:403–10.

28. Ip CLC, Loose M, Tyson JR, de Cesare M, Brown BL, Jain M, et al. MinION Analysis and Reference Consortium: Phase 1 data release and analysis. F1000Research. 2015;4:1075.

29. Hestand MS, Van Houdt J, Cristofoli F, Vermeesch JR. Polymerase specific error rates and profiles identified by single molecule sequencing. Mutat Res. 2016 Mar;784–785:39–45.

30. Garrison E, Sirén J, Novak AM, Hickey G, Eizenga JM, Dawson ET, et al. Variation graph toolkit improves read mapping by representing genetic variation in the reference. Nat Biotechnol. 2018;36:875–9.

31. Rautiainen M, Mäkinen V, Marschall T. Bit-parallel sequence-to-graph alignment. Bioinforma Oxf Engl. 2019 Mar 9;

32. Jain C, Dilthey A, Misra S, Zhang H, Aluru S. Accelerating Sequence Alignment to Graphs. bioRxiv. 2019 May 27;651638.

33. Kurtz S, Phillippy A, Delcher AL, Smoot M, Shumway M, Antonescu C, et al. Versatile and open software for comparing large genomes. Genome Biol. 2004;5:R12.

34. Šošić M, Šikić M. Edlib: a C/C ++ library for fast, exact sequence alignment using edit distance. Bioinformatics. 2017 May 1;33:1394–5.

35. Pruitt KD, Tatusova T, Maglott DR. NCBI reference sequences (RefSeq): a curated non-redundant sequence database of genomes, transcripts and proteins. Nucleic Acids Res. 2007 Jan 1;35(suppl_1):D61–5.

36. Li H, Handsaker B, Wysoker A, Fennell T, Ruan J, Homer N, et al. The Sequence Alignment/Map format and SAMtools. Bioinformatics. 2009 Aug 15;25:2078–9.

37. Ono Y, Asai K, Hamada M. PBSIM: PacBio reads simulator—toward accurate genome assembly. Bioinformatics. 2013 Jan 1;29:119–21.

