## Supplementary figures and figure texts for "Increasing the efficiency of long-read sequencing for hybrid assembly with k-mer-based multiplexing"

### Supplement:

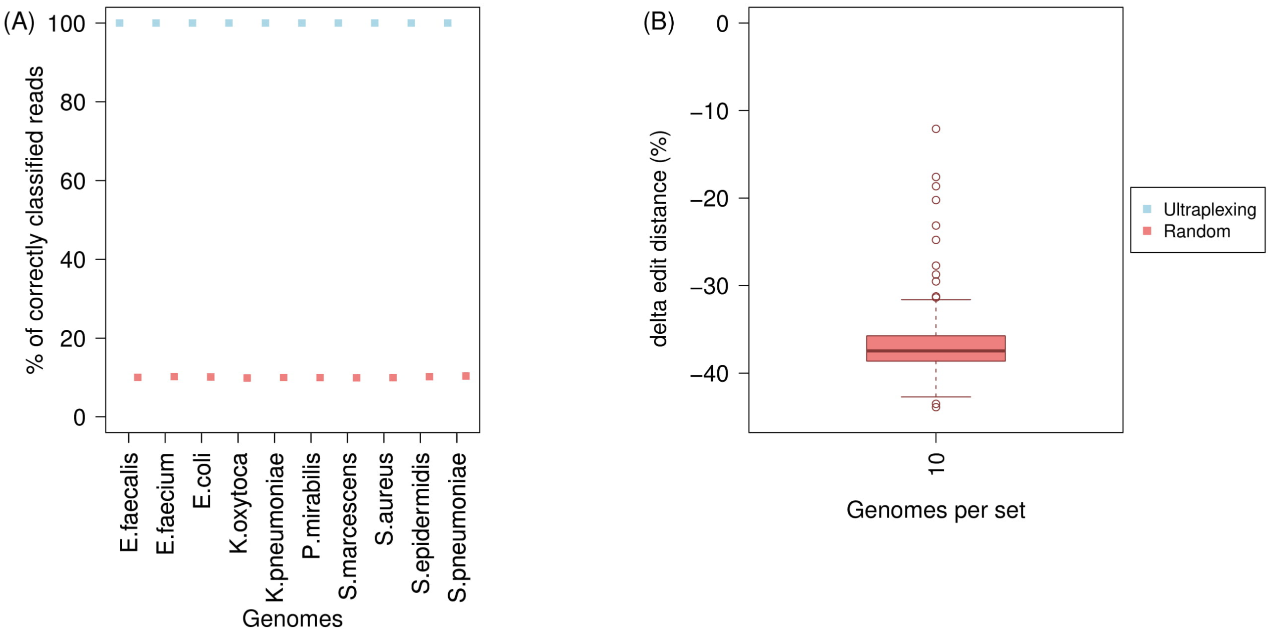


*Supplementary Figure 1: Read classification in a simulation experiment with ten different human pathogens. The figure shows the percentage of correctly classified simulated long reads (A) and Δedit distance for falsely classified reads (B). Reads were assigned according to the Ultraplexing algorithm (Ultraplexing) and randomly (Random).*

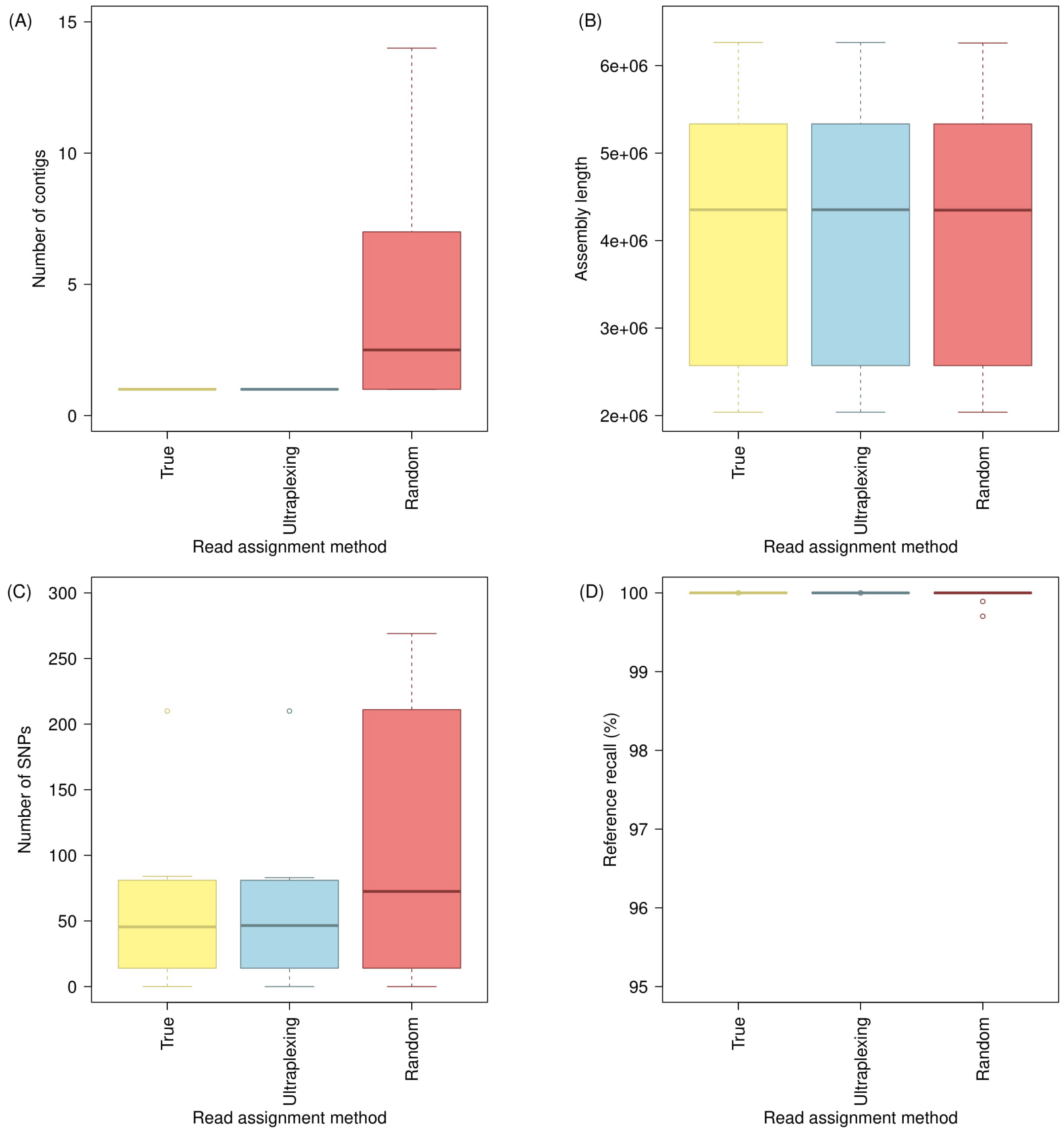


*Supplementary Figure 2: Assembly accuracy in a simulation experiment with ten different human pathogens. The figure shows the distribution of contigs per assembly (A); the distribution of assembly lengths (B); the distribution of SNPs per assembly (C); and the distribution of reference recall (D). Long reads were assigned to their true origin (True); by the Ultraplexing algorithm (Ultraplexing); and randomly (Random). Independent of long-read assignment method, the same simulated short-read data are used for all hybrid assemblies of the same species. SNPs and reference recall were calculated relative to the utilized reference genomes.*

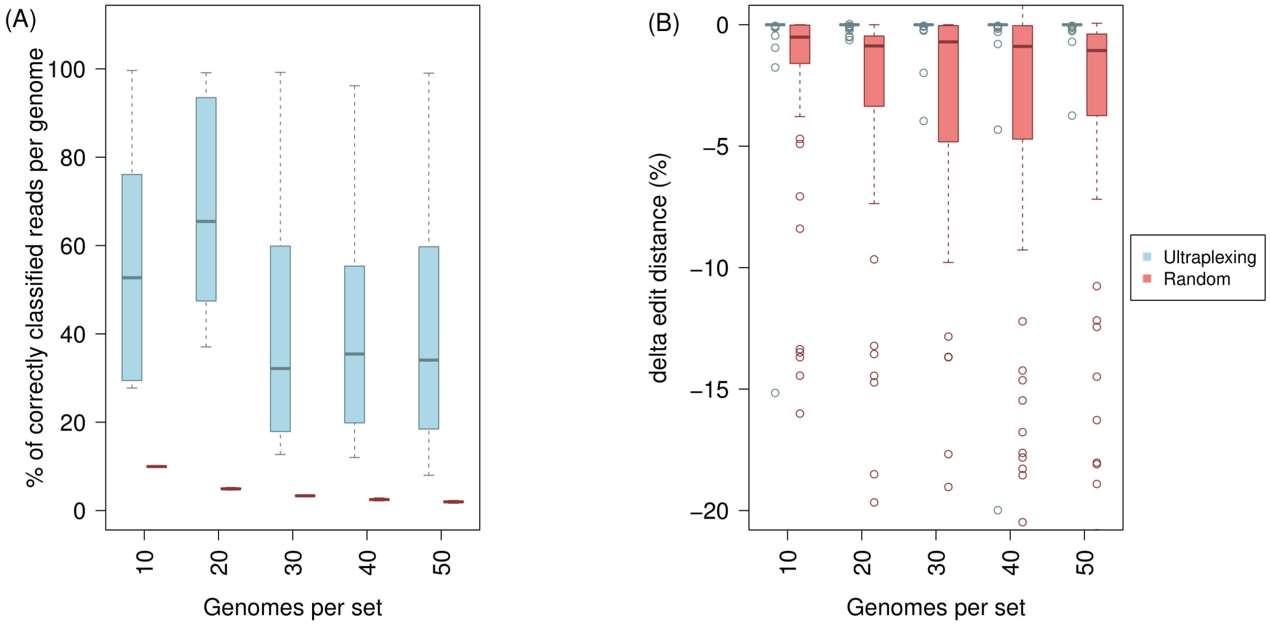


*Supplementary Figure 3: Read classification in five simulation experiments with 10 – 50 different plasmid-containing S. aureus genomes that all contain plasmids. The figure shows the distribution of the percentage of correctly classified simulated long reads (A) and the distribution of Δedit distance for falsely classified reads (B). Reads were assigned by the Ultraplexing algorithm* (Ultraplexing) *and randomly (Random).*

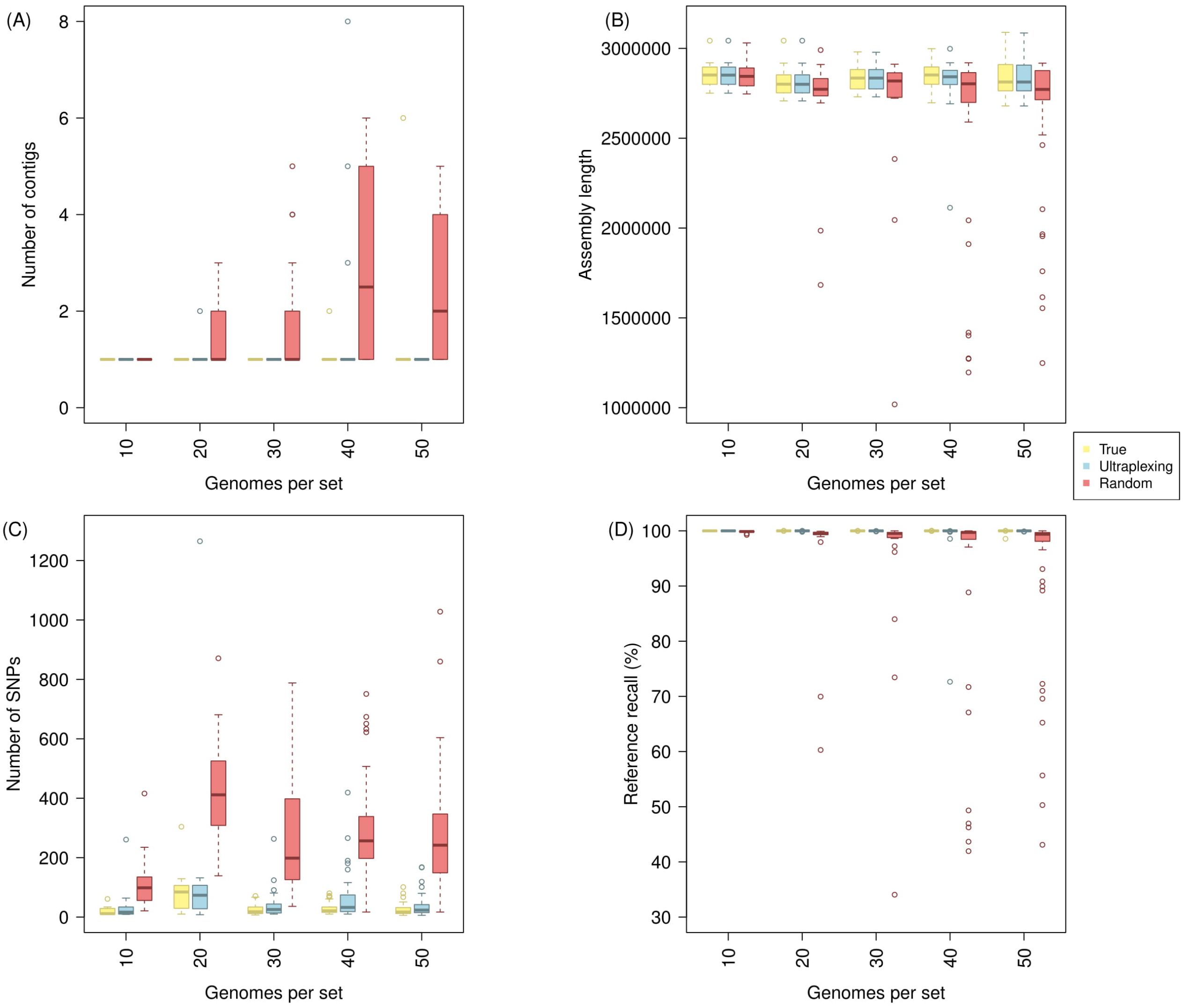


*Supplementary Figure 4: Chromosomal assembly accuracy in five simulation experiments with 10 – 50 different plasmid-containing S. aureus genomes. Reference and assembly contigs were classified as 'chromosomal' or 'plasmid' and evaluated separately (see Methods); shown here are results for the 'chromosomal' compartment. The figure shows the distribution of contigs per assembly (A); the distribution of assembly lengths (B); the distribution of SNPs per assembly (C); and the distribution of reference recall (D). Long reads were assigned to their true origin (True); by the Ultraplexing algorithm (Ultraplexing); and randomly (Random). Independent of long-read assignment method, the same simulated short-read data are used for all hybrid assemblies of the same isolate. SNPs and reference recall were calculated relative to the utilized reference genomes.*

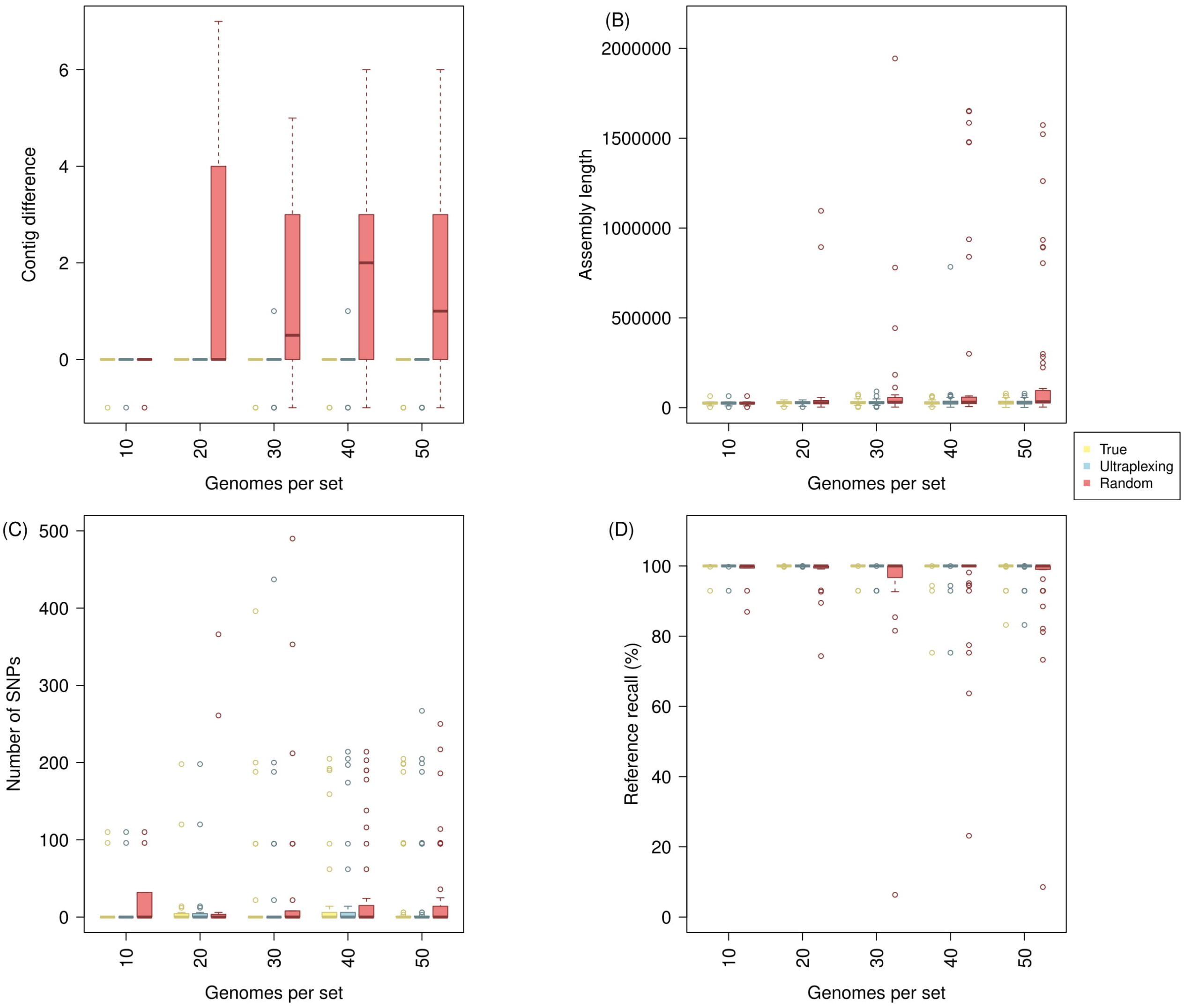


*Supplementary Figure 5: Plasmid assembly accuracy in five simulation experiments with 10 – 50 different plasmid-containing S. aureus genomes. Reference and assembly contigs were classified as 'chromosomal' or 'plasmid' and evaluated separately (see Methods); shown here are results for the 'plasmid' compartment. The figure shows the distribution of contigs per assembly (A); the distribution of assembly lengths (B); the distribution of SNPs per assembly (C); and the distribution of reference recall (D). Long reads were assigned to their true origin (True); by the Ultraplexing algorithm (Ultraplexing); and randomly (Random). Independent of long-read assignment method, the same simulated short-read data are used for all hybrid assemblies of the same isolate. SNPs and reference recall were calculated relative to the utilized reference genomes.*
