## Supplementary table texts for "Increasing the efficiency of long-read sequencing for hybrid assembly with k-mer-based multiplexing"

Supplementary Table 1:
Read classification and assembly accuracy in a simulation experiment with ten different human pathogens. The first sheet shows the number and proportion and correctly classified reads for Ultraplexing and random read assignment (Tab 1: Classifier); the second sheet shows Δedit distance data for a subsample of individual reads falsely classified by the 'random' assignment method (Tab 2: Prediction Score); the third sheet shows assembly metrics (contig lengths, assembly precision, reference recall, SNPs and N50/L50) for different long-read assignment methods (Tab 3: Assembly). Metrics with the suffix '_longest' refer to the longest contig in the corresponding assembly. 'True': long reads assigned to their true origin; 'predicted': long reads assigned by the Ultraplexing algorithm; 'random': long reads assigned randomly. Individual columns are visualized Supplementary Figures 1 and 2.

Supplementary Table 2:
Read classification and assembly accuracy in five simulation experiments with 10 – 50 different *S. aureus* genomes. The first sheet shows the number and proportion and correctly classified reads for Ultraplexing and random read assignment (Tab 1: Classifier); the second sheet shows Δedit distance data for subsamples of individual reads falsely classified (Tab 2: Prediction Score); the third sheet shows assembly metrics (contig lengths, assembly precision, reference recall, SNPs and N50/L50) for different long-read assignment methods (Tab 3: Assembly). Metrics with the suffix '_longest' refer to the longest contig in the corresponding assembly. 'True': long reads assigned to their true origin; 'predicted': long reads assigned by the Ultraplexing algorithm; 'random': long reads assigned randomly. Individual columns are visualized Figure 2.

Supplementary Table 3:
Read classification and assembly accuracy in five simulation experiments with 10 – 50 different plasmid-containing *S. aureus* genomes. The first sheet shows the number and proportion and correctly classified reads for Ultraplexing and random read assignment (Tab 1: Classifier); the second sheet shows Δedit distance data for subsamples of individual reads falsely classified (Tab 2: Prediction Score); the third sheet shows assembly metrics (contig lengths, assembly precision, reference recall, SNPs and N50/L50) for different long-read assignment methods (Tab 3: Assembly). For assembly accuracy metrics, reference and assembly contigs were classified as 'chromosomal' or 'plasmid' (see Methods) and evaluated separately (column 'Type_2'). Metrics with the suffix '_longest' refer to the longest contig in the corresponding assembly. 'True': long reads assigned to their true origin; 'predicted': long reads assigned by the Ultraplexing algorithm; 'random': long reads assigned randomly. Individual columns are visualized in Supplementary Figures 3, 4 and 5.

Supplementary Table 4:
Assembly accuracy in real-data experiments. The first sheet (Tab 1: Assembly) shows results for a multiplexed sequencing run of 10 *S. aureus* isolates (Type_2: ultraplex_10x); results for a multiplexed sequencing run of 48 different *S. aureus* isolates benchmarked against an initial set of reference genomes (Type_2: ultraplex_48x); and results for a 10-isolate subset from the 48-sample run benchmarked against an improved set of reference genomes (Type_2: ultraplex_48, with '_n' suffix in column 'Type_1'). 'Predicted': long reads assigned by the Ultraplexing algorithm; 'random': long reads assigned randomly. The second sheet (Tab 2: Reference) shows the properties of the utilized reference genomes

Supplementary Table 5:
Names and NCBI accessions of the genomes underlying the simulation experiment with ten different human pathogens (Tab1: Multi-species); the simulation experiments with 10 – 50 *S. aureus* genomes (Tab2: Multi-strain); and the simulation experiments with 10 – 50 plasmid-containing *S. aureus* genomes (Tab3: Multi-strain plus plasmids). Note that the second and third sheet specify the sets of genomes that the 10 – 50 samples were drawn from; Supplementary Table 3 and 4 also specify the accessions of the selected isolates.

Supplementary Table 6:
Incorrectly assembled incompletely recovered plasmids in the simulated sets with 10 – 50 *S. aureus* isolates. The table shows the difference in contig number between the assembly and the source genome and a description of the identified assembly error.

Supplementary Table 7:
BLAST results for contigs putatively representing plasmids in two real-data experiments, based on assemblies of molecularly barcoded data. The table shows identity and E-value of the 5 best BLAST hits for each contig. The first sheet (Tab 1: Ultraplex_10_Isolates) shows results for the first real-data experiment (sequencing of 10 *S. aureus* isolates) and the second sheet (Tab 2: Ultraplex_48_Isolates) shows results for the repeat run of the second real-data experiment (re-sequencing of 10 out of 48 *S. aureus* isolates).Shown are the 5 best hits (corresponding to identity and e-value) for each isolate.

Supplementary Table 8:
Metrics for sequencing reads assigned to individual isolates in the real-data experiment with 48 different *S. aureus* isolates, for molecularly barcoded Nanopore data (Type: original) and for the separate 48-sample Nanopore run demultiplexed by the Ultraplexing algorithm (Type: predicted).

Supplementary Table 9:
Summary statistics of all real-data read sets (Oxford Nanopore and Illumina). The metrics are partitioned into blocks for each real-data experiment (sequencing of 10 isolates, sequencing of 48 isolates, sequencing of 10 out of 48 isolates).
